# Fecal pollution explains antibiotic resistance gene abundances in anthropogenically impacted environments

**DOI:** 10.1101/341487

**Authors:** Karkman Antti, Pärnänen Katariina, Larsson D.G. Joakim

**Affiliations:** Department of Infectious Diseases, Institute of Biomedicine, The Sahlgrenska Academy, University of Gothenburg, Guldhedsgatan 10, SE-413 46, Gothenburg, Sweden; Center for Antibiotic Resistance research (CARe) at University of Gothenburg, P.O. Box 440, SE-40530, Gothenburg, Sweden; Faculty of Biological and Environmental Sciences, University of Helsinki, Viikinkaari 1, 0014 University of Helsinki, Finland; Department of Microbiology, University of Helsinki, Viikinkaari 9, 00014 University of Helsinki, Finland

## Abstract

Discharge of treated sewage leads to release of antibiotic resistant bacteria, resistance genes and antibiotic residues to the environment. Such pollution can directly contribute to increased morbidity caused by the transmission of resistant fecal pathogens. Residual antibiotics in wastewaters have been speculated to select for resistant bacteria and thereby promote the evolution and emergence of new resistance factors. Increased abundance of antibiotic resistance genes in sewage and sewage-impacted environments may, however, simply be a result of fecal contamination with resistant bacteria rather than caused by an on-site selection pressure. In this study we have disentangled these two alternative scenarios by relating the relative resistance gene abundance to the accompanying extent of fecal pollution in publicly available metagenomic data. This was possible by analyzing the abundance of a newly discovered phage which is exceptionally abundant in, and specific to, human feces. The presence of resistance genes could largely be explained by fecal pollution, with no clear signs of selection in the environment, the only exception being environments polluted by very high levels of antibiotics from manufacturing where selection is evident. Our results demonstrate the necessity to take in to account the fecal pollution levels to avoid making erroneous assumptions regarding environmental selection of antibiotic resistance. The presence or absence of selection pressure has major implications for what the risk scenarios are (transmission versus evolution) and for what mitigations (reducing pathogenic bacteria or selective agents) should be prioritized to reduce health risks related to antibiotic resistance in the environment.

## Introduction

Growing concern over the threat posed by antibiotic resistant bacteria to human health has turned attention also to the environmental dimensions of the problem. Only fairly recently, the role of the environment as a source and dissemination route for antibiotic resistance has been acknowledged (Bengtsson-Palme et al., 2018; Berendonk et al., 2015; Wright, 2010). Treated sewage effluent discharge is one of the most important point sources of resistant bacteria and resistance genes release to the environment (Karkman et al., 2018; Rizzo et al., 2013). Along with gut microbes, which contain a wide array of resistance determinants (van Schaik, 2015), antibiotics consumed by humans and animals are released into the environment in urine and fecal material contained in treated effluents from waste water treatment plants (WWTPs) and sludge applied to land.

Waste water treatment plants appear to provide suitable conditions for antibiotic resistance emergence and dissemination in bacterial populations. The incoming sewage contains bacteria from human, animal and environmental origin and a mixture of sub-therapeutic concentrations of antibiotics and other co-selective agents (Karkman et al., 2018). The WWTPs provide conditions that favor growth and metabolic activity of certain bacteria which are in close proximity to each other enabling the exchange of genetic material. For these reasons WWTPs, have been considered hotspots for antibiotic resistance emergence and dissemination (Guo et al., 2017; Rizzo et al., 2013). However, modern conventional WWTPs are generally quite effective in removing antibiotic resistant bacteria (ARB) and resistance genes (ARGs) from the raw sewage (Bengtsson-Palme et al., 2016; Karkman et al., 2016; Yang et al., 2014). While total bacterial counts are strongly reduced (often 10–1000 -fold reduction between raw sewage and treated effluents) the relative abundance to 16S rRNA gene for most ARGs also decreases or remains on similar levels. In combination, this shows that WWTPs often reduce ARB and ARG discharges efficiently (Laht et al., 2014). Despite this, sewage effluents are important point sources for resistant bacteria and resistance gene to the environment due to the large volumes released. The receiving environments form another possible hotspot for antibiotic resistance dissemination when bacteria originating from sewage and fecal material come to contact with environmental bacteria (Bengtsson-Palme et al., 2018). The fate and persistence of antibiotic resistance genes (ARGs) after they have reached the environment is not well understood.

Although a distance-decay-effect can be found at effluent release sites, the selection or dissemination of the genes has rarely been explicitly investigated (Chu et al., 2017; Czekalski et al., 2014). It has been speculated that ARGs could be selected in the receiving environment by antibiotics and other co-selective agents originating from the WWTP, contributing to further dissemination of ARGs to environmental bacteria (Berendonk et al., 2015; Eckert et al., 2018; Li et al., 2018; Martinez, 2009). However, there are experimental studies (using complex microbial communities) suggesting that the concentrations of selective agents in a sewage-impacted environment might not be sufficient to cause selection (Kraupner et al., 2018; Lundström et al., 2016) despite that competition experiments with two strains occasionally suggest otherwise (reviewed in (Andersson and Hughes, 2014)). Thus, it is equally likely that the levels of ARGs are elevated in sewage-impacted environments because of input of treated effluent containing already resistant bacteria carrying the ARGs in question. If there is selection for antibiotic resistance either in WWTPs or receiving environments, it could promote the emergence of novel resistance mechanisms and the dissemination of existing ARGs. Selection pressure would facilitate novel resistance mechanisms and the maintenance of recently transferred ARGs in the new host (Bengtsson-Palme et al., 2018). Antibiotic resistance genes are often associated with mobile genetic elements (MGEs) making them more prone to being transferred between bacterial species (Stokes and Gillings, 2011) and some antibiotics are known to promote horizontal transfer of MGEs (Roberts and Mullany, 2009). Even in the absence of selection, plasmids can transfer to new hosts with high frequency (Hall et al., 2017), but for the plasmids to be maintained in the new host, selection for some trait encoded in that plasmid is often needed. MGEs are abundant in sewage effluents and elevated levels have been detected in effluent receiving environments (Guo et al., 2017), but their detailed dynamics are largely unknown, as with ARGs.

To determine the resistance levels in WWTPs and receiving environments several methods have been used including selective culturing of indicator bacteria, quantitative PCR (qPCR) of resistance genes and metagenomics (Bengtsson-Palme et al., 2016; Harris et al., 2014; Karkman et al., 2016; Laht et al., 2014; Sidrach-Cardona et al., 2014; Yang et al., 2014). The class 1 integron (CL1) integrase gene has often been used as a proxy for anthropogenic impact and total antibiotic resistance gene abundance with good resolution (Gillings et al., 2015). However, as CL1s can contain a wide array of resistance genes and, thus, can be subjected to selection themselves, they are not an independent measure which can be used to assess the selection or dissemination of ARGs. Fecal pollution levels have rarely been incorporated in the determination of possible section or dissemination of resistance genes. This is likely mainly because many of the fecal markers are not easy to quantify. Detecting fecal marker bacteria using metagenomics is often difficult due to the low abundance of common marker bacteria in the whole community (Bengtsson-Palme et al., 2017). A robust marker for fecal pollution would provide the means for distinguishing between on-site selection and dissemination of the genes versus accumulation and to observe decrease in ARGs which is due to dilution of fecal pollution in the receiving environments. Therefore, taking fecal pollution into account in models of ARB and ARG pollution from sewage would help in determining the critical control points for antibiotic resistance selection by enabling distinguishing between accumulation and selection.

Recently, crAssphage, a bacteriophage most probably infecting the genera *Bacteroides* and *Prevotella*, was identified from human fecal metagenomes. It is an abundant human intestinal phage, as approximately 1.7 % of human fecal metagenome reads align to it, and it is six times more abundant in public metagenomes compared to all the other know phages together (Dutilh et al., 2014). While highly abundant in human feces, it is considerably rarer in feces from other animals (García-Aljaro et al., 2017). The high abundance of crAssphage in human feces as well as its host specificity makes it valuable for microbial source tracking. Indeed, it has been shown to perform equally well or better than traditional markers in qPCR assays (Stachler et al., 2017). Given its high abundance, it can be used in metagenomic studies for estimating the human fecal pollution levels in the environment (Ahmed et al., 2018a; Stachler and Bibby, 2014). crAssphage has been shown to correlate moderately with only a few ARGs in storm drain outfalls (Ahmed et al., 2018b) and thus, could be a good candidate for estimating the effect of fecal pollution in ARG dynamics. Moreover, it is not physically linked to resistance genes like integrons are and hence likely independent from ARGs and thus, could enable the determination of ARG horizontal gene transfer and selection patterns in receiving environments. Another fecal *Bacteroides* phage, ϕB124-14, has been used for microbial source tracking but unlike crAssphage, it is also abundant in porcine and bovine gut (Ogilvie et al., 2017), therefore making it possibly a good candidate for estimating the combined effect of human and production animal fecal pollution.

In this study, we have quantified the abundance of acquired ARGs and fecal pollution levels using a selected set of public metagenomes from sewage polluted environments and, in addition, nearly 500 metagenomes from MG-RAST. We show that in practically all the studied environments, the ARG levels correlate with fecal pollution levels with no signs of selection or dissemination of the resistance genes, except in sediments polluted with wastewater from drug manufacturing containing exceptionally high levels of antibiotics (Bengtsson-Palme et al., 2014; Kristiansson et al., 2011), where ARG abundance did not correlate with the fecal pollution, showing clear signs of selection. Fecal pollution also correlated well with the CL1 and MGE abundance in receiving environments. Using fecal samples from (Hu et al., 2013) and the Integrative Human Microbiome project (iHMP) (Integrative HMP (iHMP) Research Network Consortium, 2014), we also show that although both the acquired mobile ARGs and the crAssphage are abundant in human feces, their abundance is not correlated in the fecal metagenomes confirming the independence of these two measures. On the other hand, we found a significant but weak correlation between the ARG abundance and CL1. Taken together, this approach could be used in future studies or re-analyzing already available samples in order to assess and detect selection and/or dissemination patterns of ARGs in environments with sewage pollution.

## Materials and methods

### Data collection

#### Phage genomes

The crAssphage (NC_024711.1) and ϕB124-14 (HE608841.1) genomes were downloaded from GenBank and indexed for mapping using bowtie2-build (Langmead and Salzberg, 2012).

#### Studies selected

Metagenomic studies on human impacted environments were searched from the literature and six studies where the sequencing data was available were selected and downloaded from either SRA or ENA (Bengtsson-Palme et al., 2014; Chu et al., 2017; Kristiansson et al., 2011; Lekunberri et al., 2018; Marathe et al., 2017; Ng et al., 2017; Rowe et al., 2016, 2017). The studies included samples from river and lake sediments, WWTP and hospital effluents and river water. Two waste water treatment plant studies with comprehensive metadata were used to determine the impact of waste water treatment on the ARG abundance and fecal content (Bengtsson-Palme et al., 2016; Petrovich et al., 2018). Accession numbers for all metagenomic data can be found from Supplementary Table 1. Many data sets from peer-reviewed studies that were candidates for being included in this study were unfortunately not made publicly available.

#### MG-RAST

A set of 484 metagenomes from MG-RAST, excluding metagenomes from different human body sites, analyzed in (Pal et al., 2016), were used for the study. The metagenomes included soil, freshwater, marine, animal, waste water, agricultural and air samples. Full list of accessions and annotations are available in Supplementary Table 1

#### Human gut metagenomes

To study the abundance of crAssphage and its association to the total ARG abundance in human gut metagenome samples, 74 Chinese and 234 European subjects were used from a previous study (Hu et al., 2013). In addition, gut metagenomes from 141 US subjects were downloaded from the HMP portal (https://portal.hmpdacc.org/). Accessions and download links for HMP subjects can be found from Supplementary Table 1.

#### Animal gut metagenomes

To determine the abundance of crAssphage in animal gut metagenomes, 12 chicken (Xiong et al., 2018), a subset of 100 samples from a bigger pig metagenome study (Xiao et al., 2016) and 42 cow rumen metagenomes (Stewart et al., 2018) were downloaded from SRA. Accession numbers are in Supplementary Table 1.

### *crAssphage* and ARG annotation

Metagenomic reads were mapped against the crAssphage genome using bowtie2 (Langmead and Salzberg, 2012) and the crAssphage genome coverage was calculated using Samtools (Li et al., 2009). Only read pairs mapping in proper pairs were calculated in case of paired-end sequencing. For single end metagenomes, the mapped reads were filtered with quality value 10 using Samtools. The genome coverage was used as a measure for crAssphage abundance in the sample. ResFinder, a database of mobile, acquired antibiotic resistance genes (Zankari et al., 2012) was translated into amino acid sequences using Biopython. The longest ORF was selected for each gene. MGE database from Pärnänen et al. (2018) (*in review*) (available at: https://github.com/KatariinaParnanen/MobileGeneticElementDatabase) was used for MGE annotation by first removing all plasmid marker sequences. All remaining entries were translated to amino acid sequence as described earlier. ARGs and MGEs were annotated against the translated ResFinder/MGE database using DIAMOND blastx (Buchfink et al., 2015) with the following parameters: minimum identity 90 %, minimum match length 20 AA. In case of paired-end sequencing, matches on the second read were counted only if there was no match on the first read. Both crAssphage abundance and antibiotic resistance gene abundance were normalized with the total base pair count in the metagenome. Mapping against ϕB124-14 was done exactly as with crAssphage. All source code for the data analysis can be found from https://github.com/karkman/crassphage_project.

### Statistical analyses

Both normalized crAssphage abundance and normalized total ARG abundance were log10 transformed and linear regression was done using the *lm* function in R v.3.2 (R Core Team, 2016). The ARG abundances in the MG-RAST metagenomes were predicted from the crAssphage abundance based on the linear model based on the selected studies using the function *predict* in R. Figures were drawn in R using the base graphic package and ggplot2 package v.2.2.1 (Wickham, 2009). Smoothing curves using a linear model were drawn with function *geom_smooth* in ggplot2. Statistical comparison of mobile antibiotic resistance gene abundance between European, Chinese and US subjects was done using the *aov* function in R. A *post-hoc* test was done using the *TukeyHSD* function in R. All R code for the data analysis can be found from https://github.com/karkman/crassphage_project.

## Results and discussion

Concerns about the elevated levels of antibiotic resistant bacteria and resistance genes in the receiving environments of treated sewage have been raised in several publications (Berendonk et al., 2015; Eckert et al., 2018; Martinez, 2008, 2009; Rowe et al., 2017). However, based on only abundance data it is difficult to determine whether the increase is explained by selection and dissemination to the resident microbiota or by constant input of fecal bacteria. Therefore, we analyzed mobile antibiotic resistance genes in metagenomes from environments with anthropogenic impact from sewage discharge and correlated the total abundance of the mobile ARGs with a human fecal pollution marker, crAssphage, abundance. Our results show that the observed ARG abundances strongly correlate with crAssphage, meaning that the fecal pollution levels largely explain the observed abundances and there are now clear signs of wide scale selection or dissemination of antibiotic resistance in the affected environments.

### Screening of human fecal metagenomes to evaluate the suitability of crAssphage in assessing antibiotic resistance gene dynamics

To determine the independence of crAssphage and mobile ARGs in human feces, we analyzed fecal metagenomes from 74 Chinese, 234 European (Hu et al., 2013) and 141 subjects from the USA (HMP) (Suppl. Table 1) for ARG and crAssphage abundance. The screening of fecal metagenomes proved that crAssphage had a potential for revealing selection dynamics in receiving environments by correlating ARG abundance with the phage abundance in environments with human fecal pollution. We did not find any correlation between the ARG and crAssphage abundance in fecal metagenomes of the studied populations (linear regression, F = 25.51, adj. R^2^ = 0.21, p > 0.05, Figure 1, Suppl. Table 2), confirming that ARG abundance is independent from crAssphage abundance. On the other hand, the correlation between total ARG abundance and *intI1* gene abundance was significant when taking in to account the different base levels of ARG abundance in the populations (linear regression, F = 20.55, adj. R^2^ = 0.29, p < 0.05, Figure 1, Suppl Table 2) showing the expected dependence between ARGs and CL1s.

**Figure 1.**
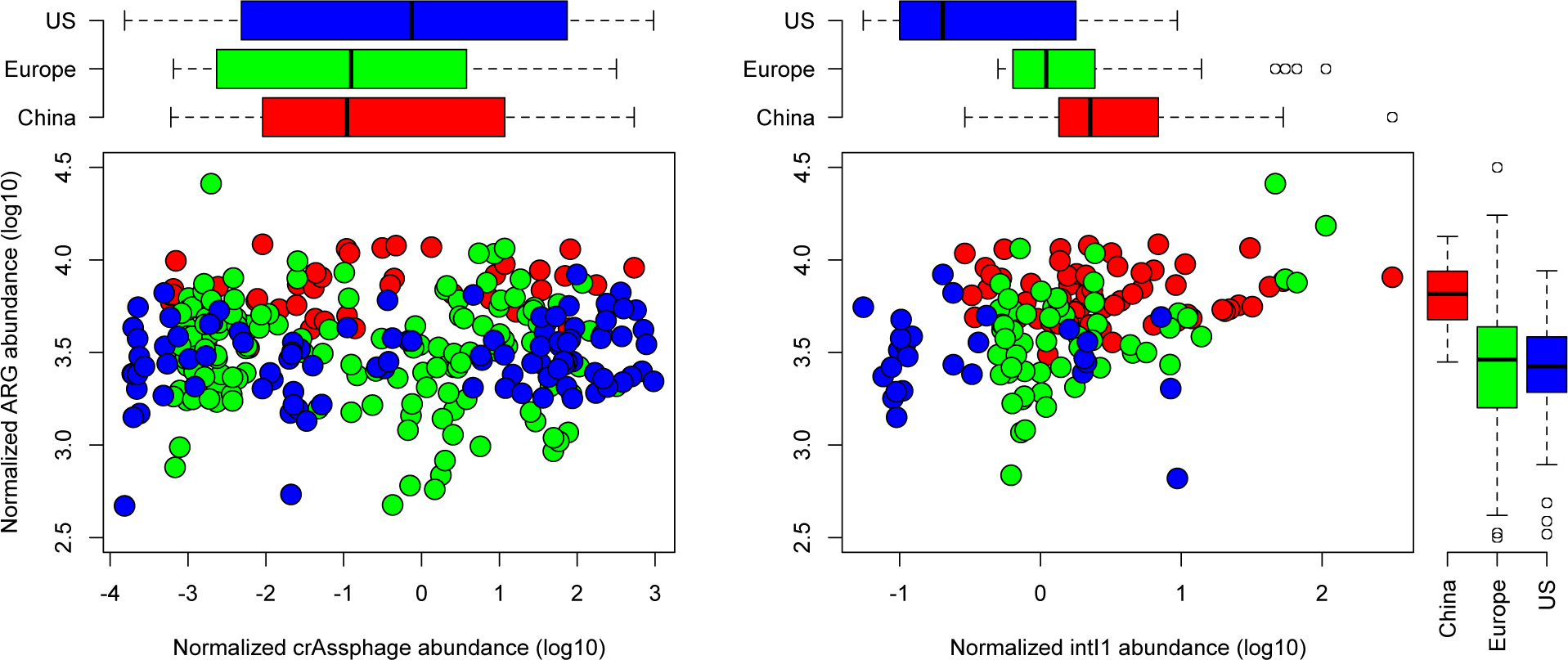
Mobile antibiotic resistance gene abundance in relation to crAssphage (left) and *intI1* gene (right) abundance in studied human fecal metagenomes from (Hu et al., 2013) and Integrative Human Microbiome Project (iHMP). We found no correlation between the total mobile ARG abundance and crAssphage abundance in human fecal metagenomes confirming the independence of the abundance of mobile ARGs from crAssphage abundance.

We observed that the crAssphage abundance was similar in all cohorts, even though the abundance varied dramatically between individual subjects (Figure 1). The uniform abundance of crAssphage across the studied populations suggests that crAssphage could be used as a fecal pollution marker globally, which is in line with previous observations (Cinek et al., 2018). However, it should be noted that in another study, crAssphage was reported to be less abundant in sewage from Asia and Africa compared to Europe and US (Stachler and Bibby, 2014). In our analysis, the total relative ARG abundances were similar in US and European subjects, while the Chinese subjects had higher relative abundance of resistance genes in their fecal metagenomes (Analysis of variance, Tukey *post-hoc* test, adj. p < 0.05 for both, Figure 1). The higher ARG levels in Chinese subjects compared to Europeans, using the same samples which were analyzed in here, has been reported earlier (Hu et al., 2013). Differences on the ARG levels on population level might originate from the historical amount of antibiotics used regionally as well as the degree of transmission control. However, even with differences on the initial crAssphage to ARG ratio in different populations, deviations from this relationship in the receiving environment will reveal possible selection hotspots.

### Correlation of fecal pollution with antibiotic resistance gene abundance in anthropogenically impacted environments reveal a hotspot for selection in industrially polluted sediments

To determine the relationship between ARG abundance and fecal pollution in environments with anthropogenic impact, we selected metagenomic studies from literature where sequence data was available (Suppl. Table 1). In all environments, except in environments polluted directly by waste water from the manufacturing of antibiotics (see below), the total ARG abundance positively correlated with the crAssphage abundance, showing that in these environments the ARG abundance could largely be explained by the extent of fecal pollution (linear regression, F = 32.51, adj. R^2^ = 0.75, p<0.05, Figure 2, Suppl. Table 2). The highest ARG abundances were detected in industrially polluted Indian sediments and in hospital and WWTP effluents from UK and Singapore. Lower levels of ARGs were detected in sediments and river water downstream of a hospital as well as WWTP effluent discharge points, in line with the diluted fecal material quantified with crAssphage abundance (Suppl. Table 2).

**Figure 2.**
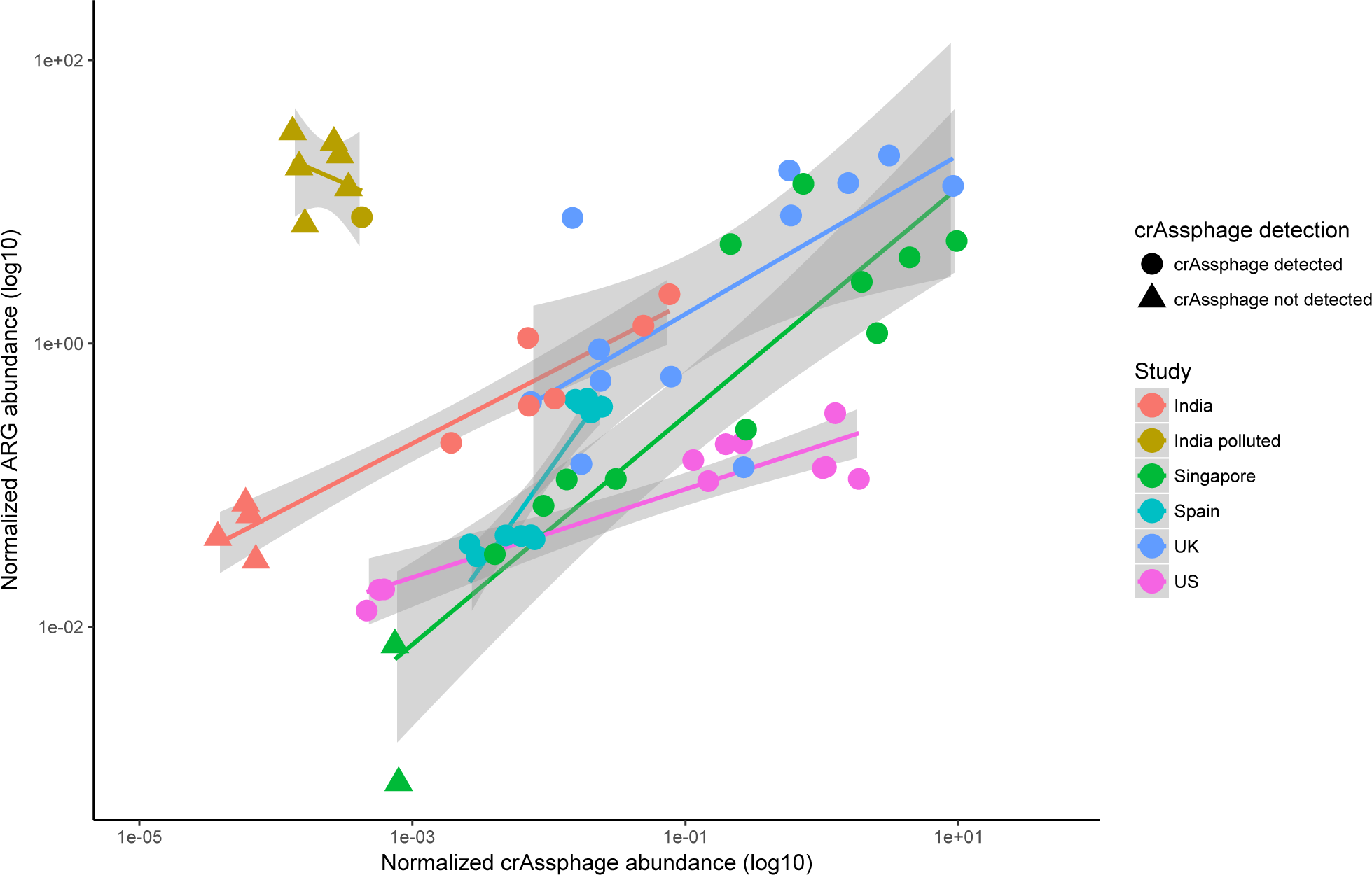
Correlation between ARG abundance and crAssphage abundance in environments with pollution from WWTPs, hospitals or drug manufacturing. The environments effected by drug manufacturing are polluted with exceptionally high levels of antibiotics, and the analyses show clear selection for antibiotic resistance as the ARG abundance cannot be explained by fecal pollution. ARG abundance and crAssphage abundance were normalized with total nucleotide count in the metagenomes. For samples where we did not detect crAssphage (indicated by triangles), half of the detection limit (corresponding to one read mapping to crAssphage) was used and normalized to the total nucleotide count.

Antibiotic resistance genes conferring resistance to different classes of antibiotics followed the same trend in all datasets (Suppl. figure 1), with the exception for quinolones in two datasets. The deviations, however, were explained by a single gene, *oqxB*, coding for part of an RND efflux pump with very broad substrate specificity (Hansen et al., 2007). Also, the counterpart of the mobilized RND system, *oqxA* did not follow the pattern of *oqxB*. This suggests that the elevated presence of reads matching *oqxB* in some samples reflects a taxonomic change towards bacteria carrying the gene chromosomally, and not likely as a consequence of quinolone exposure and selection by quinolones.

Overall, the strong and consistent correlation between ARGs and crAssphage suggests that the observed enriched ARG abundance is primarily due to fecal contamination rather than selection of the ARGs in the downstream environments. Similarly, the ARG richness, total abundance of MGEs and *intI1* gene abundance could also be explained by fecal pollution levels (Suppl. Figure 2).

There is a clear exception with regards to Indian sediments polluted with exceptionally high levels of antibiotics from drug manufacturing, where ARG levels were very high, at the same time crAssphage was not detected in most of these samples indicating they contained very little human fecal material. These and other sediments from the same industrially polluted area contain the highest levels of antibiotics ever measured in the environment. Together with the exceptionally high abundance of ARGs and very few or no crAssphage reads in the metagenomes provides strong support for direct selection rather than fecal contamination (Kristiansson et al., 2011). It should be noted that the DNA in these samples were amplified using RepliG prior to sequencing, which could favor small circular plasmids and thereby inflate the counts for genes carried by these (Bengtsson-Palme et al., 2014). However, even by removing the most abundant ARGs, the pattern was still consistent (Suppl. Figure 3).

### Large scale analysis of metagenomes shows that antibiotic resistance gene abundance is largely explained by fecal pollution rather than selection

To expand our analysis beyond the selected studies of polluted environments, we analyzed 484 publicly available metagenomes from MG-RAST, previously analyzed for ARGs in Pal et al. (2016). We were able to detect crAssphage only in samples taken from the Mississippi river (USA), WWTPs, activated sludge with high ammonia content, Beijing air and from laboratory mice. Fecal pollution correlated with the observed ARG abundance in the river water and Beijing air and in waste waters excluding activated sludge with high ammonia content (Figure 3). The strong correlation of ARGs and crAssphage in the large scale analysis of metagenomes are in line with our results in the selected anthropogenically polluted sites, confirming that also in these environments the mobile ARG abundance could be explained with fecal pollution and no apparent signs of large-scale selection or horizontal gene dissemination could be detected (Figure 3, Suppl. Table 2). Interestingly, this was true also for Beijing smog samples, hosting a particular high diversity of ARGs (Pal et al., 2016). The activated sludge samples with high ammonia concentrations were from a laboratory experiment (Crovadore et al., 2017) and conclusions about possible selection during waste water treatment in these samples should not be drawn.

**Figure 3.**
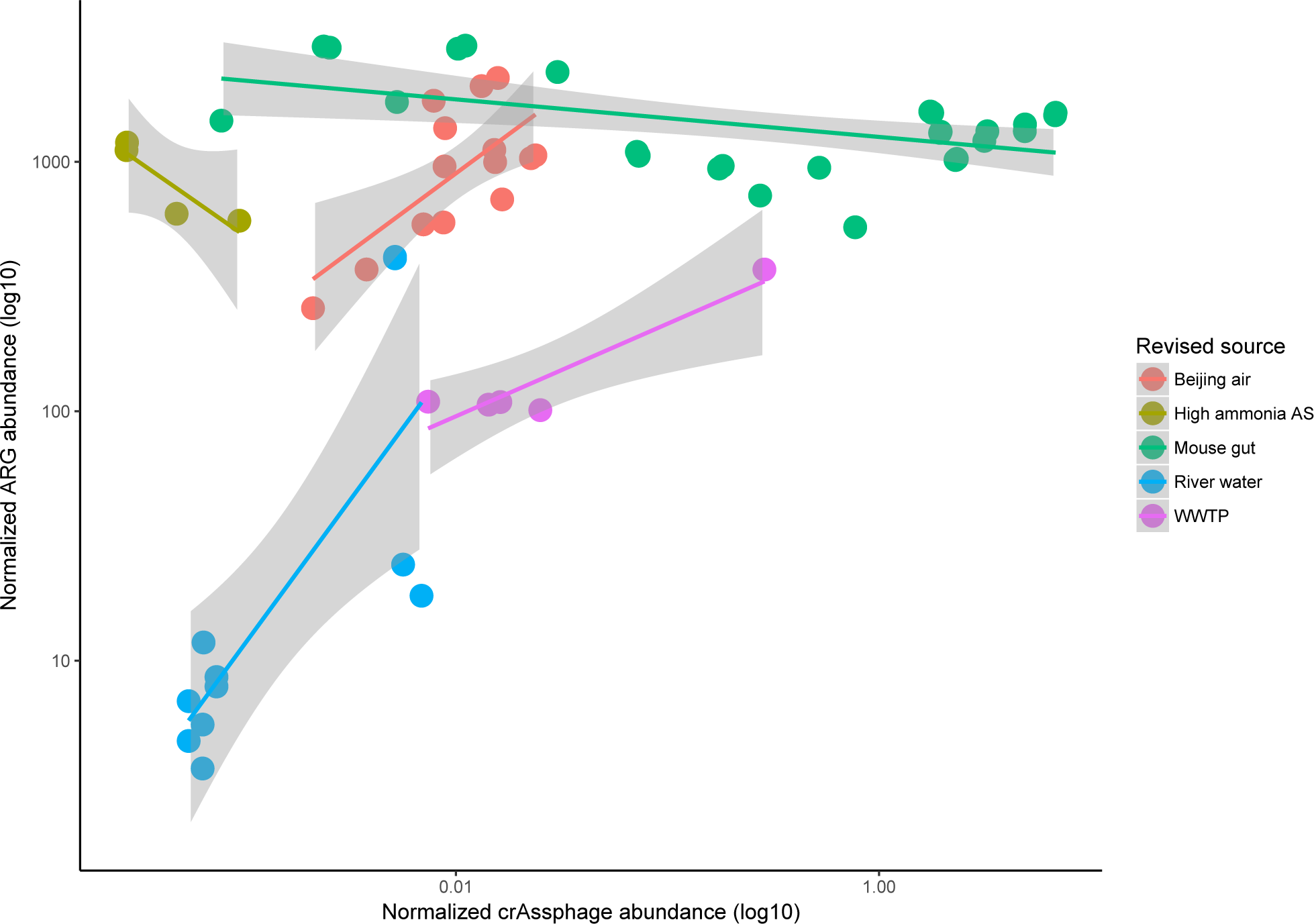
The correlation between crAssphage abundance and total ARG abundance in MG-RAST metagenomes where crAssphage was detected. Only in the mice gut and sludge with high ammonia concentration we could not see a correlation (discussed in main text). Original MG-RAST feature annotations were revised manually using the project and sample descriptions due to the misleading information in original annotations. Original annotations can be seen in Suppl. Table 1.

A sometimes high abundance of crAssphage and consistently high abundances of ARGs were found from mice gut metagenomes in MG-RAST. All of these mice were from murine model experiments and were not given antibiotics (Everard et al., 2014; Langille et al., 2014; Rooks et al., 2014). It is known that laboratory mice gut microbiota bear some similarities to human gut microbiota (Nguyen et al., 2015) and differs from the gut microbiota of wild mice (Kreisinger et al., 2014; Rosshart et al., 2017), which could explain these findings. As in human feces, the crAssphage abundance in mice feces was not linked to the total ARG abundance.

The samples with elevated resistance levels where we could not detect crAssphage were either polluted with fecal material from non-human sources or were fecal metagenomes from other animals than humans (Suppl. Figure 4). We were not expecting crAssphage to correlate with ARG abundance in environments with non-human fecal pollution. To confirm this we analyzed 12 chicken gut (Xiong et al., 2018), 100 pig gut (Xiao et al., 2016) and 42 cow rumen (Stewart et al., 2018) metagenomes and did not detect crAssphage. This, on the other hand, was expected as crAssphage is far less abundant in other animals and in sites polluted with other than human fecal material (Ahmed et al., 2018a; Dutilh et al., 2014; García-Aljaro et al., 2017; Stachler and Bibby, 2014) making it specific for detecting anthropogenic pollution.

To test whether another fecal marker would work for environments polluted with other than human fecal material, we also determined the abundance of ϕB124-14 phage in the MG-RAST metagenomes. We found it to be less abundant than crAssphage in human impacted environments and not to perform well with other pollution sources and did not use it in any further analyses (data not shown).

### Predicting antibiotic resistance gene abundance with crAssphage

Using a linear model constructed from the selected metagenomes from polluted environments, we were able to predict the differences in antibiotic resistance gene abundance in the environments with anthropogenic impact in MG-RAST (Beijing air, river water & WWTP) with the crAssphage abundance with good accuracy (linear regression, F = 34.76, adj. R^2^ = 0.54, p < 0.05, Suppl. Figure 5). However, due to the different base levels of resistance genes in the point source (sewage), the baseline (intercept) of ARG abundance cannot be predicted with high accuracy. It has been shown that different parts of the world have different resistance burden (Hu et al., 2013), which is seen also in fecal samples analyzed in this study (Figure 1). Furthermore, different environments might have different levels of fecal pollution from domestic animals, which could explain some of the varying background levels of resistance when compared to the crAssphage abundance. However, when the baseline level of an environment is estimated, the model using crAssphage performs well in predicting the ARG abundance.

### Analysis of selection for antibiotic resistance genes in waste water treatment plants

As noted earlier, WWTPs could potentially serve as hotspots for antibiotic resistance selection and horizontal gene transfer. To determine correlation between ARG abundance and fecal pollution in WWTPs during the treatment process, we analyzed metagenomes from three Swedish WWTPs (Bengtsson-Palme et al., 2016) and two WWTPs in Wisconsin, USA (Petrovich et al., 2018). These studies were selected since the sequence data was publicly available and was accompanied with comprehensive metadata from the WWTPs sampled. The results show clearly the ARG abundance decreases from raw sewage to the treated sewage with decreasing fecal material during the treatment process (Figure 4). In sludge, the ARG abundance seems to be even lower than expected based on the crAssphage abundance (Suppl. Figure 6). These results speak against WWTPs being hotspots with a strong antibiotic resistance selection acting as a driver for horizontal gene transfer (Guo et al., 2017; Rizzo et al., 2013) and show that at least in the studied treatment plants the treatment process eliminates ARGs along with fecal pollution with high efficiency. Also other studies have shown the reduction of ARGs during the treatment process (Karkman et al., 2016; Laht et al., 2014) and our results connect the reduction to elimination of fecal material from sewage. In all of the five treatment plants studied, the correlation between crAssphage and ARG abundance was similar (Figure 4). A linear model between ARG abundance and crAssphage detection confirmed that the correlation was significant and similar in both countries and all WWTPs (linear regression, F = 71.26, adj. R^2^ = 0.59, p<0.05). Only the intercepts differed between the Swedish and USA WWTPs indicating in this case that the proportion of ARGs to crAssphage in the sewage entering the plant was different in the two countries, but the ARG removal efficiencies of the treatment processes were on par for all treatment plants.

**Figure 4.**
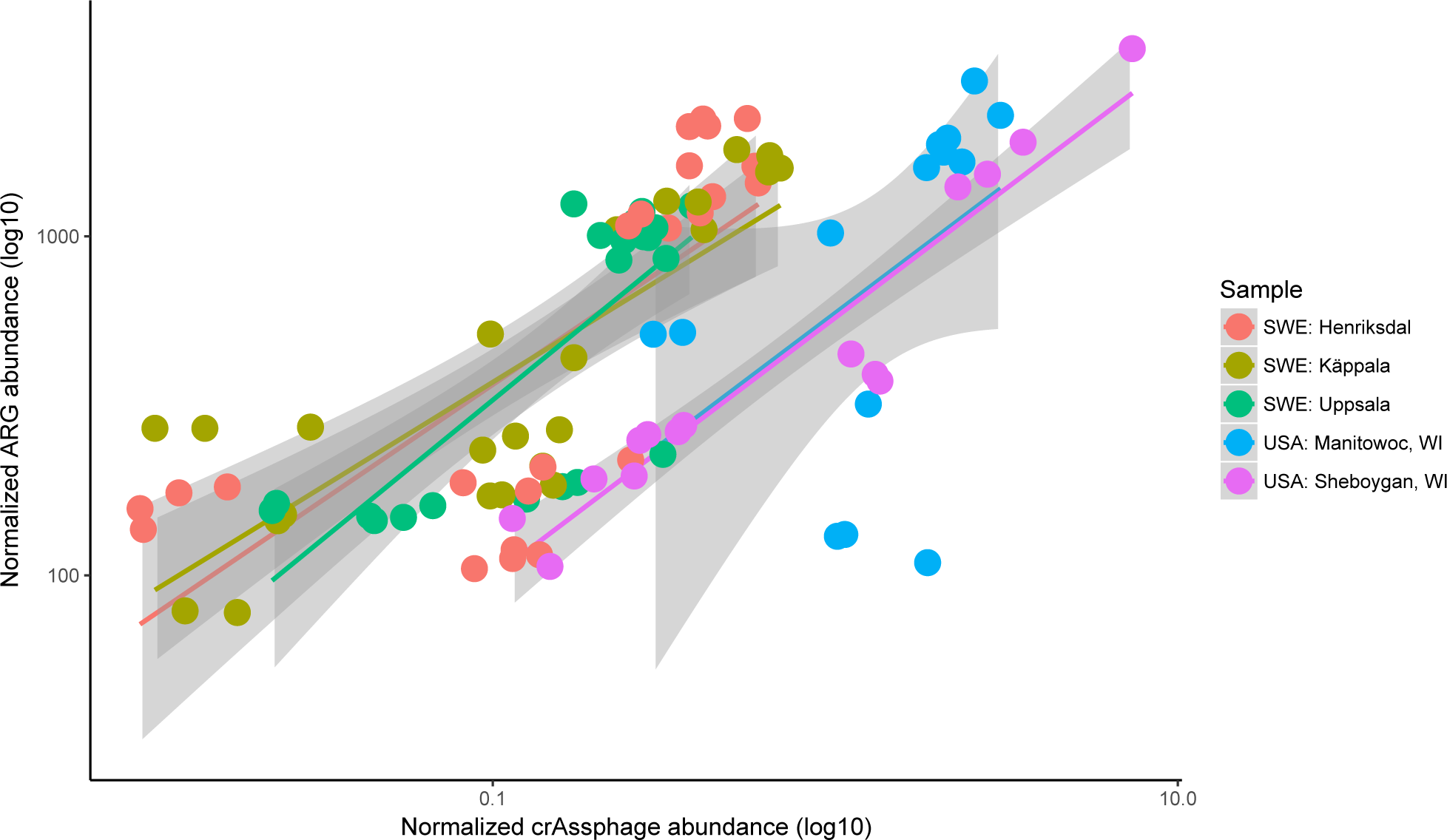
ARG abundance and crAssphage detection in two US and three Swedish waste water treatment plants showing similar correlation with different base level of resistance. Smoothing curves based on linear model separately for each plant are shown with 95 % confidence intervals in grey.

One would perhaps expect a more similar ARG/crAssphage ratio in Swedish and US sewage influents, given the overall similar resistance gene abundance in gut metagenomes of Swedish and US subjects (Feng et al., 2018). Explanations behind the differences found might be technical and come from sample handling or be related to the sources of sewage entering the plant but unfortunately, we cannot asses that with the data available. The ARGs per DNA read were, on average, slightly higher and crAssphage more common in the US populations (refer to Figure 1 to see differences in ARG abundances in different countries). This suggest that one should probably restrict comparisons of ARG/crAssphage ratios in samples that are contaminated by the same population of people and prepared using similar protocols, as we have done in this study.

### Estimated resistance risk correlates with fecal pollution levels in metagenomic samples with anthropogenic impact

A computational pipeline for estimating the risk of resistance from metagenomic sequence data in a given environment was published recently (Oh et al., 2018). We analyzed the same samples that were used for benchmarking the pipeline and found that the resistance risk followed fecal pollution levels in the environments with human fecal pollution and the correlation was even stronger with the total count of mobile ARGs in all environments (Suppl. Figure 7). Our results show that in one of the hospital samples there was elevated levels of resistance compared to the fecal content (Suppl. Figure 7). However, their overall contribution to the resistance load at WWTPs has been shown to be small (Buelow et al., 2018). So, to fine-tune the risk assessment, we would propose to include the fecal content in to the analysis to be able to determine possible selection or dissemination scenarios.

Estimating the risk associated with environmental antibiotic resistance is far from simple. Besides the total amount of ARGs, it clear that association of resistance genes with MGEs and pathogens elevates the risk caused to human health (Martínez et al., 2014). However, detecting a resistance gene in the environment does not necessarily mean a risk for human health. Certainly, there is a higher probability of transfer when there are more transferrable genes and recipient cells together and when the genes are already on mobile genetic elements ready to be transferred to pathogens. However, selection plays an important role in the processes required for the transferred gene to persist in the new host or a newly emerged gene to disseminate (Bengtsson-Palme and Larsson, 2015). Using proxies for fecal pollution such as crAssphage enable detecting selection and thus, can help in assessing risks associated with it.

## Conclusions

Our results provide a framework for disentangling transmission of resistant, human fecal bacteria from the possible selection and horizontal gene transfer of resistance genes in the environment. We were able to detect true hotspots for antibiotic resistance gene selection in sediments receiving exceptionally high levels of antibiotics from industry. In addition, we show that in all other studied environments receiving anthropogenic waste, there was no evidence of wide scale selection. On the contrary, the ARG abundance correlated strongly with fecal pollution, which does not support the prevailing speculations that major selection for antibiotic resistance occurs in WWTPs or effluent receiving environments. A lack of apparent selection in these environments means that the emergence of new resistance determinants is less likely and the transferred resistance genes are not likely to be fixated on new hosts due to selection pressure. In the heavily polluted Indian sediments new resistance determinants are possibly more persistent and disseminate more efficiently in the population due to the competitive advantage they give compared to the receiving strains. It should be noted that selection occurring on small scale, which does not affect the entire population, or selection of limited types of resistance genes would probably not be detected using this method. Furthermore, rare horizontal transfer events will definitely be missed and more precise methods are needed.

Another important source of antibiotic resistance determinants is animal husbandry, which uses more antibiotics than are prescribed to humans. In the US animal manure exceeds the human sewage sludge by approximately 100-fold (Gerba and Smith, 2005). As crAssphage is abundant only in human fecal material it cannot be used to estimate fecal content of environments receiving animal manure and feces. However, it is likely that similar phages that are specific to different production animals could be identified and used in a similar fashion. Discovering more makers for fecal pollution advance the field of studying antibiotic resistance in the environment. We argue that the use of crAssphage, or similar fecal pollution markers, should be incorporated broadly in studies determining the selection, persistence and fate of resistance genes originating from fecal pollution, which is likely the main source of resistance genes of environmental ARG pollution.

In terms of ranking risk in resistomes, we argue that the total count of ARGs or their genetic context does not give the whole picture of the actual risks. Without a way to estimate the extent of selection pressure in the environment, the total count of ARGs or the association of an ARG to mobile element adds only a few pieces to the entire puzzle of disentangling the risks related to resistance genes, especially when in most cases the genes are eventually diluted to near zero concentrations following the diminishing fecal pollution. Real-time PCR primers for detecting crAssphage are already available (Stachler et al., 2017), so the use of metagenomics is not necessary, making the analysis quick, easy and affordable. When collecting data, we encountered many studies where sequencing data was not available, even though sometimes presented as available for download in public repositories. Re-analyzing these samples using markers for fecal pollution would expand our knowledge on resistance gene dynamics is diverse environments. To conclude, we argue that including a proxy for fecal pollution in future studies would enable a more comprehensive understanding of the antibiotic resistance dynamics in the receiving environments, providing more reliable estimates of the risk scenarios and perhaps most importantly, discovering true environmental hotspots for selection.

